# The perceptual and spatial architecture of Müllerian mimicry in *Heliconius* Butterflies

**DOI:** 10.64898/2026.03.02.709095

**Authors:** Christopher Lawrence, Michelle Ramirez, Tanya Berger-Wolf, Owen McMilan, Daniel Rubenstein

## Abstract

Müllerian mimicry theory predicts reciprocal convergence among defended species toward shared warning signals, yet the extent, symmetry, and perceptual basis of this convergence remain poorly quantified. Using deep-learning–based phenotyping combined with neural networks calibrated to avian predator and butterfly visual systems, we quantify mimetic similarity across 56 subspecies of *Heliconius erato* and *H. melpomene*. Mimetic phenotypes show broad clustering consistent with recognized mimicry rings, but vary continuously within. Phenotypic similarity is weakly structured by phylogeny and strongly organized geographically, consistent with repeated local convergence. Spatial overlap among co-mimics is heterogeneous, suggesting that frequency-dependent selection operates at variable spatial scales across mimicry communities. Models parameterized to different visual systems reveal that trait salience and discriminatory performance depend on the observer, indicating that apparent imperfect mimicry may arise from receiver-specific perceptual weighting. Asymmetric learning between lineages further suggests that convergence may not always be reciprocal, consistent with one species evolving towards another in advergence. Together, these results recast mimicry as a perceptually structured and spatially dynamic continuum rather than a set of discrete, mutually reinforcing rings, highlighting how community composition and sensory ecology shape the evolution of adaptive resemblance.

## Background

Mimicry has played a foundational role in evolutionary theory, providing a model for how natural selection shapes adaptive phenotypic diversity^1^. Among Müllerian systems, *Heliconius* butterflies are emblematic: across the Neotropics, chemically defended species share similar warning color patterns organized geographically into distinct “mimicry rings”^2^. Classical theory predicts that positive frequency-dependent selection drives convergence toward a locally shared warning signal, reducing phenotypic diversity within communities ^3,4^. Yet *Heliconius* exhibits striking spatial variation in colour pattern^5^, and mimetic resemblance among co-occurring species is rarely exact (Figure 1)^6–8^. This diversity raises questions about the strength, precision, and symmetry of mimicry convergence, and about how mimicry is structured across space.

**Figure 1.**
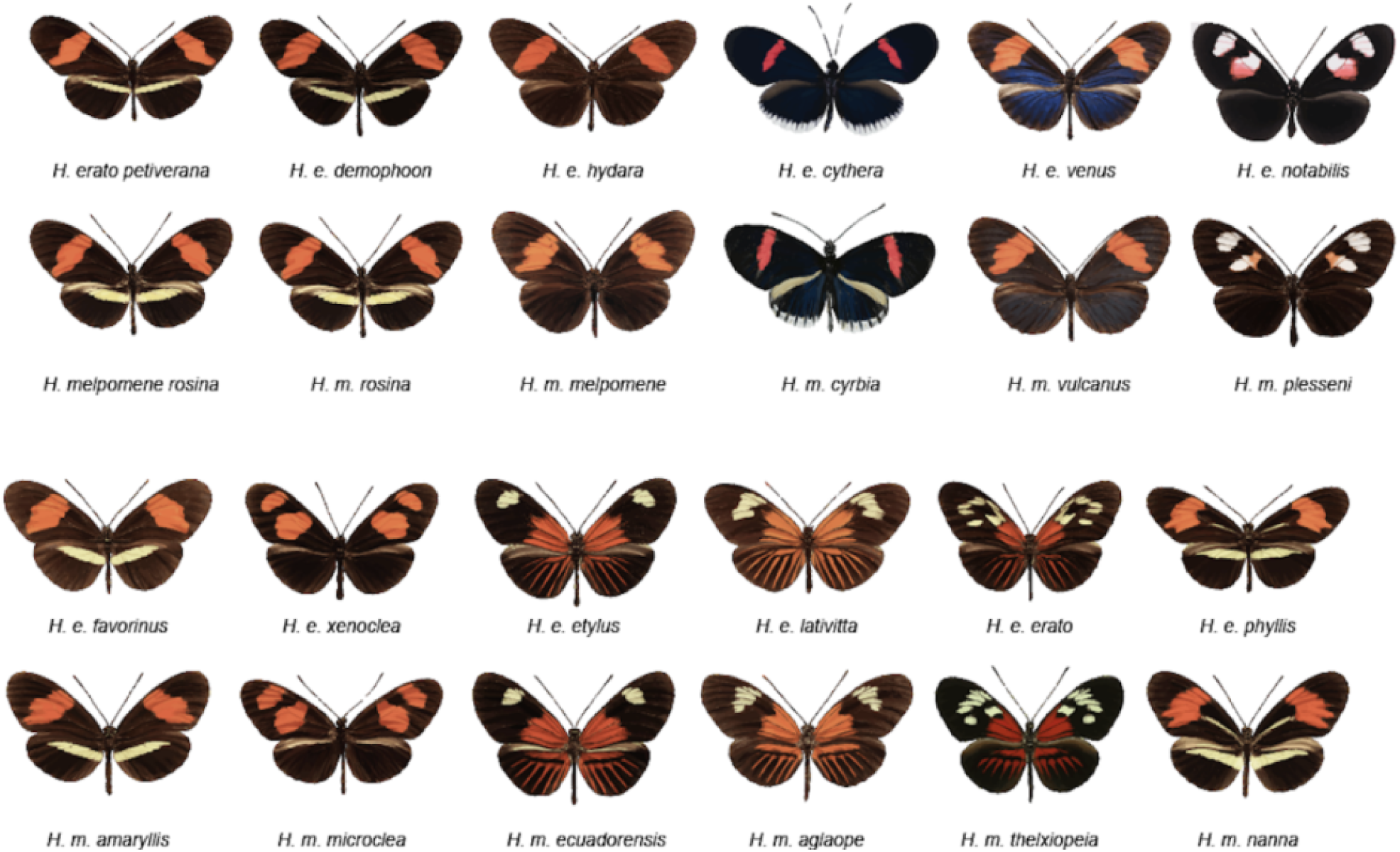
Mimetic convergence and divergence in *Heliconius erato* and *H. melpomene*. Representative dorsal images of co-mimetic subspecies illustrating convergence on shared warning color patterns across lineages and divergence in specific pattern elements among and within mimicry rings.

A central limitation in addressing these questions is that mimetic relationships are typically treated as categorical ^9^. Species are assigned to mimicry rings based on shared motifs, but resemblance itself is rarely quantified^10^. As a result, mimicry is often described as present or absent, and convergence is assumed to be reciprocal ^11^. Without continuous measures of similarity, it remains difficult to evaluate how strongly co-mimics converge, whether convergence is symmetric, how it is structured by geography and ancestry, and whether apparent “imperfections” are perceptually meaningful to predators.

Here we reconstruct the architecture of Müllerian mimicry in *Heliconius* using continuous measures of perceptual distance derived from biologically calibrated deep learning models. By quantifying resemblance within a perceptual framework, we test whether mimicry forms discrete rings or continuous gradients, whether similarity reflects shared ancestry or repeated convergence, how this similarity is organized geographically, and whether convergence between co-mimetic lineages is reciprocal. We further model alternative visual acuity regimes to determine how observer identity shapes perceived similarity and which pattern elements most strongly mediate signal discrimination.

We show that mimetic phenotypes broadly correspond to recognized rings but vary continuously within rather than forming discrete categories. Similarity is only weakly predicted by phylogenetic relatedness, yet strongly structured by geography, indicating repeated convergence shaped by regional selective environments. Spatial overlap among co-mimics is heterogeneous, suggesting that the scale and symmetry of frequency-dependent selection differ across communities. Models calibrated to alternative visual systems further reveal that the traits defining mimicry depend on the observer, rendering apparent imperfections perceptually contingent. Together, these results indicate that *Heliconius* mimicry is not a fixed set of reciprocally defined rings, but a geographically structured and receiver-dependent process in which resemblance operates along a continuum.

### Measuring Mimetic Fidelity

We focused on *Heliconius erato* and *H. melpomene* because their mimicry is widely described as approximately one-to-one, providing a natural test of the strength and symmetry of Müllerian convergence. Using standardized dorsal images of 56 subspecies (27 *H. erato* and 29 *H. melpomene*), we quantified resemblance with a pre-trained convolutional neural network (CNN) validated for modeling perceptual similarity^12^. This approach yields continuous distances that approximate discrimination judgments, allowing mimicry to be measured rather than assigned categorically.

Subspecies assigned to the same mimicry ring clustered in perceptual space (Fig. 2a), forming two primary groups corresponding to the “postman” and “rayed” mimicry rings, blue and yellow respectively. However, similarity among members of the same ring varied quantitatively. For example, phenotypes with double red fore-wing bands grouped more closely with other red-banded forms than with double white-banded forms assigned to the rayed ring. This pattern indicates that color similarity contributes more strongly to mimetic resemblance than band number alone, suggesting that specific visual traits disproportionately structure mimicry within rings.

**Figure 2.**
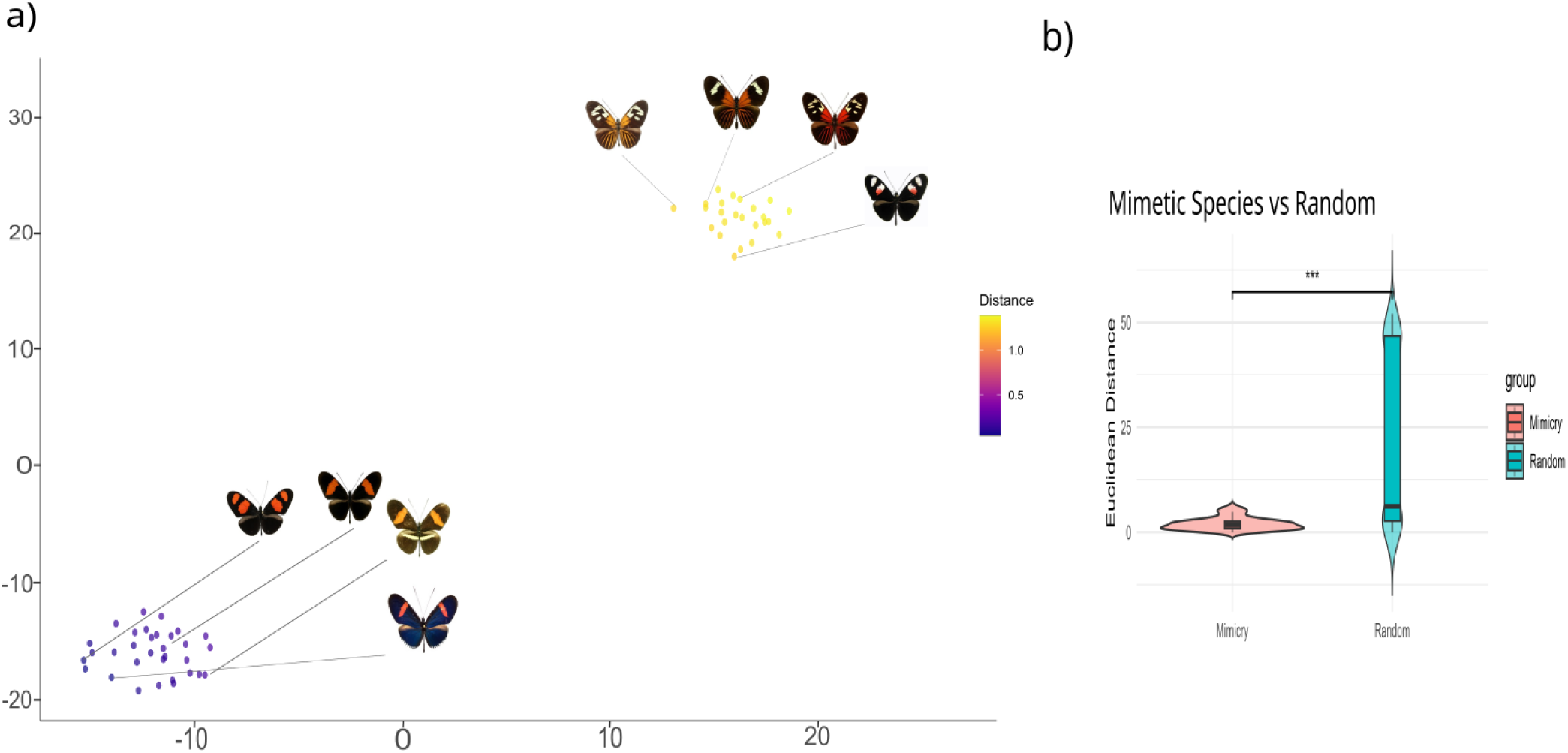
Quantification of mimetic color patterns in *Heliconius erato* and *Heliconius melpomene*. *(a)* t-distributed stochastic neighbor embedding (t-SNE) plot visualizing the perceptual embedding of mimetic *Heliconius erato* and *Heliconius melpomene* subspecies. *(b)* Violin plot comparing the Euclidean distances in perceptual space between known mimetic pairs (red) and 1,000 randomly generated species pairs (blue).

Perceptual distances also recovered historically recognized model–mimic subspecies pairs, which were significantly more similar than randomly assembled pairs (*p* < 0.001; Fig. 2b). Together, these results show that mimicry can be quantified as a continuous trait that both recovers established Müllerian relationships and reveals variation within and among mimicry rings.

### Mapping Mimicry on a phylogeny

To determine whether mimetic similarity is constrained by shared ancestry (phylogenetic inertia) or instead reflects repeated adaptive convergence, we explicitly tested for phylogenetic signal in warning pattern phenotype ^13^. Because these analyses require a resolved phylogeny, we restricted this test to 44 species (of the original 56) for which reliable phylogenetic information was available^14,15^. If phenotype is constrained by evolutionary history, closely related taxa should resemble one another, and phenotypic similarity should mirror phylogenetic relatedness.

Conversely, weak phylogenetic signal would suggest evolutionary lability and repeated convergence across lineages. A distance-based phenotypic tree showed substantial topological incongruence with the species phylogeny (Fig. 3a), indicating that closely related taxa do not consistently share similar warning patterns. We quantified phylogenetic signal using Blomberg’s K^16^ and Pagel’s *λ*^17^. Blomberg’s K indicated weak and non-significant signal (K = 0.0365, p = 0.193), whereas Pagel’s *λ* yielded a moderate but significant estimate (*λ* = 0.52, p = 0.02). These results suggest that evolutionary history exerts a partial constraint on warning pattern evolution. Mimetic resemblance is therefore only partially phylogenetically structured, consistent with repeated convergent evolution across lineages.

**Figure 3.**
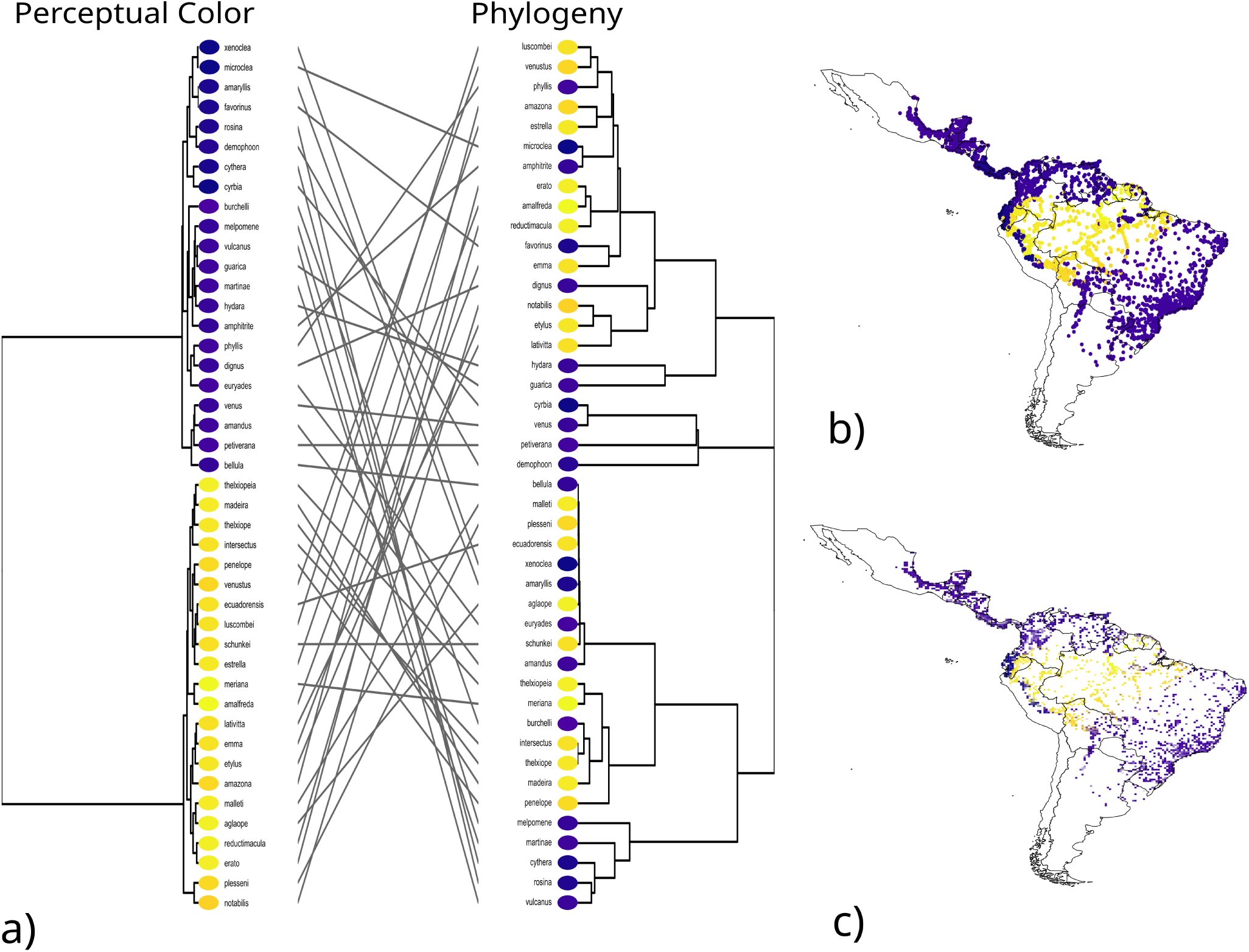
Discordance between phylogeny and perceptual similarity, and geographic structure of mimicry in *Heliconius* butterflies. *(a)* Tanglegram comparing the topology of color pattern similarity (left) and phylogenetic relationships (right) across mimetic *Heliconius erato* and *Heliconius melpomene,* color-coded according to its perceptual embedding. *(b)* High-resolution geographic map illustrating the spatial distribution of perceptual color pattern similarity across Central and South America. *(c)* Map of average perceptual color embedding scores across the geographic range.

### Geographic Variation of Mimicry

We next asked whether geographic proximity better predicts mimetic similarity than shared ancestry (Fig. 3a). Subspecies-level phenotypic distances were integrated with georeferenced occurrence data^18^, allowing resemblance to be visualized across the Neotropics. Clear regional clustering emerged (Fig. 3b), with spatial groupings corresponding to established mimicry rings. Amazonian populations were dominated by yellow-shifted phenotypes, whereas Andean and northern South American regions exhibited cooler-toned clusters. A spatially smoothed surface further revealed gradients and transition zones aligned with known Müllerian ring boundaries and regions of elevated phenotypic diversity (Fig. 3c). In contrast to the limited phylogenetic structuring observed above, mimetic similarity was strongly organized by geography at a continental scale. These results indicate that convergence in *Heliconius* is structured primarily by regional selective environments and community context rather than by shared ancestry.

### Quantifying Mimicry Overlap

To test whether co-mimetic taxa co-occur geographically—as predicted under positive frequency-dependent selection—we quantified spatial niche overlap among established mimicry pairs. Under Müllerian theory, predators select against rare phenotypes, which should favor geographic concordance among species sharing a warning signal^19^. Using kernel density estimates from ∼16,065 occurrence records, we calculated Schoener’s *D* for 16 mimetic pairs.

Overlap values ranged from 0.148 to 0.998 (mean = 0.789), indicating generally high but heterogeneous geographic congruence (Figure 4). Several pairs exhibited near-complete overlap (e.g., *H. erato microclea* and *H. melpomene xenoclea*, *D* > 0.98), consistent with strong spatial coupling under frequency-dependent selection. In contrast, other pairs sharing similar phenotypes showed only moderate or low overlap (*D* = 0.652 to 0.148), particularly within certain postman and Amazonian rayed forms. Thus, while many co-mimetic taxa co-occur extensively, the spatial coupling between them is not uniform across mimicry rings. This heterogeneity suggests that the geographic scale at which frequency-dependent selection operates may differ among communities, and that shared phenotypes can persist even where pairwise overlap is limited.

**Figure 4.**
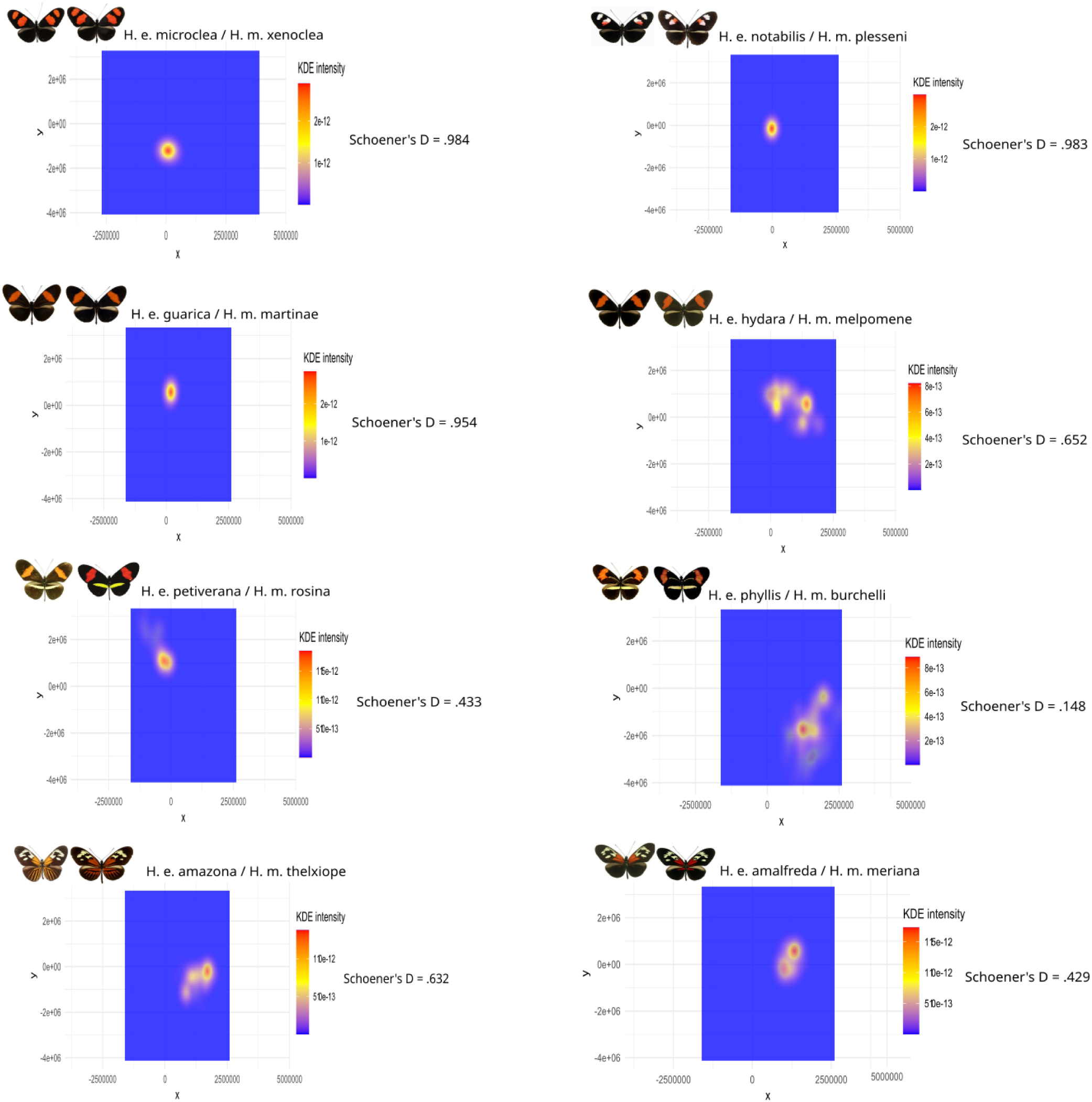
Geographic niche overlap among *Heliconius erato and Heliconius melpomene* mimetic pairs. Schoener’s D values calculated using occurrence records from *<u>Heliconius-maps.github.io</u>*. Each panel displays a pairwise comparison between mimetic taxa using kernel density heatmaps, where warmer colors indicate higher geographic overlap.

### Modeling Mimicry

Building on previous machine-learning analyses of mimicry^11^, we extended this framework to incorporate biologically calibrated visual systems. Because *Heliconius* mimicry rings involve many interacting species rather than simple model–mimic pairs, we trained models to distinguish among species while quantifying resemblance between co-mimics^10^. To examine lineage-specific versus cross-lineage recognition, we implemented three training regimes: **Eratonet**, trained exclusively on *H. erato* subspecies; **Melpomenenet**, trained exclusively on *H. melpomene* subspecies; and **Allnet**, trained jointly on subspecies from both lineages. We then simulated avian predator and butterfly visual acuity to examine how observer identity shapes mimetic discrimination^20–22^.(Figure 5a)

**Figure 5.**
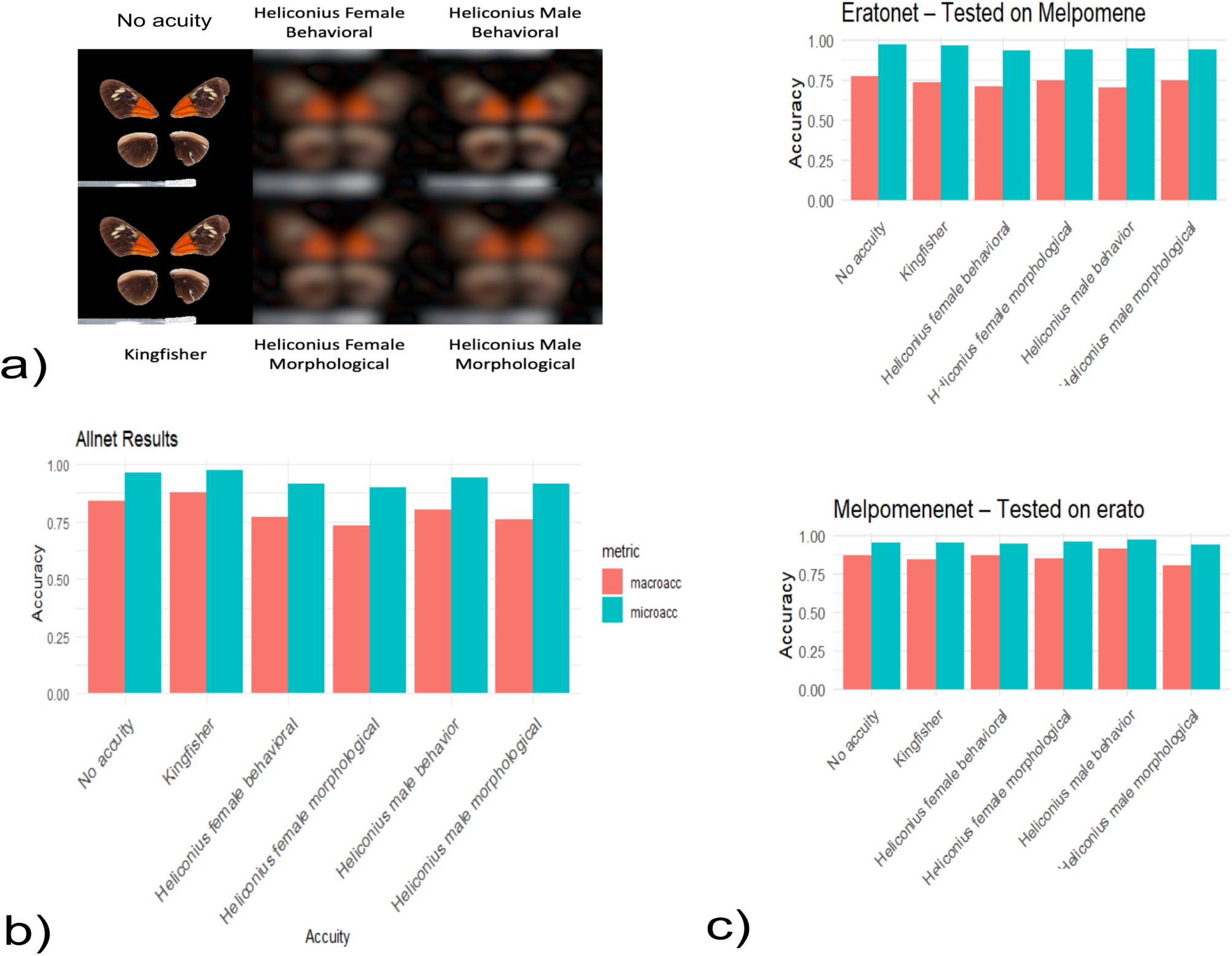
Perceptual discrimination under alternative visual acuity regimes. **(a)** Dorsal wing patterns shown under simulated avian (Kingfisher) and butterfly (male and female *Heliconius*) visual acuity. Reduced acuity smooths fine-scale pattern elements. **(b)** Classification accuracy of networks trained jointly on both lineages (Allnet) under each visual regime. Bars show microaccuracy (overall classification rate) and macroaccuracy (mean per-class accuracy), illustrating reduced perceptual separability under lower spatial resolution. **(c)** Performance of lineage-specific networks (Eratonet and Melpomenenet) under each visual regime, showing within-lineage classification and cross-lineage generalization. Cross-lineage performance differs between training directions, indicating asymmetric transfer of learned warning patterns.

We treated classification accuracy as a measure of perceptual separability: high accuracy indicates that species remain distinguishable within a given visual regime, whereas reduced accuracy reflects greater overlap in perceptual space. Using 3,822 dorsal-view images of 20 subspecies from the Cambridge Butterfly Collection^23^, we trained models under multiple acuity regimes (Figure 5a). Because *H. erato* was more densely sampled than *H. melpomene*, we report both microaccuracy (overall classification rate across all images) and macroaccuracy (mean per-class accuracy), the latter weighting each lineage equally and therefore providing a more balanced estimate under class imbalance.

Across models trained jointly on all species (Allnet), discrimination was highest under avian-calibrated vision and declined under butterfly-calibrated vision (Tigure 5b). This pattern was evident in both micro and macroaccuracy, indicating that higher spatial resolution enhances the ability to distinguish among mimetic phenotypes from different lineages. Notably, male-based visual models consistently outperformed female-based models, particularly in macroaccuracy, consistent with documented sexual dimorphism in ommatidia number and the importance of visual cues in mate recognition^24^. Reductions in acuity disproportionately affected macroaccuracy, suggesting that information loss primarily compromises discrimination among less-represented or phenotypically subtle forms.

Classical Müllerian theory assumes that co-mimetic species converge reciprocally on a shared warning signal^3^, such that learning one lineage should facilitate recognition of the other to a similar degree^25,26^. To test this prediction, we evaluated whether recognition transfers symmetrically between *H. erato* and *H. melpomene* using lineage-specific networks (Eratonet and Melpomenenet). Across all visual conditions—including predator-calibrated (Kingfisher) vision—cross-lineage generalization was asymmetric. Patterns learned from *H. melpomene* (Melpomenenet) generalized to *H. erato* more effectively than patterns learned from *H. erato* generalized to *H. melpomene*, a difference most evident in macroaccuracy (Figure 5c).This directional asymmetry indicates that cross-lineage generalization is not equivalent between lineages. The persistence of this pattern across acuity regimes—including predator-relevant visual space—suggests that perceptual overlap between lineages is structured rather than symmetric.

The highest within-lineage microaccuracy (0.983) occurred in butterfly-calibrated models trained and tested on *H. erato* (see Supplemental Table S3). Because reduced acuity removes high-frequency pattern detail, it likely compresses intra-subspecific variation, tightening class clusters and inflating microaccuracy without increasing discriminatory power. This effect was confined to within-lineage classification and did not improve cross-lineage generalization, indicating that spatial smoothing does not enhance perceptual separability among mimetic forms. Together, these results show that discriminatory performance depends not only on spatial resolution but also on taxonomic scale and lineage-specific phenotypic variance, and that recognition transfer remains asymmetric despite resolution shifts

### Identifying Mimetic Traits

To identify the traits underlying mimetic discrimination, we applied INTR^27^, a computer vision approach that generates attention heatmaps highlighting the image regions most influential for model classification. In these maps, warmer colors indicate greater contribution to recognition. Across visual conditions, models identified biologically meaningful differences, including variation in forewing band shape and hindwing banding between *H. melpomene malleti* and *H. erato lativitta* (Fig. 6a).

**Figure 6.**
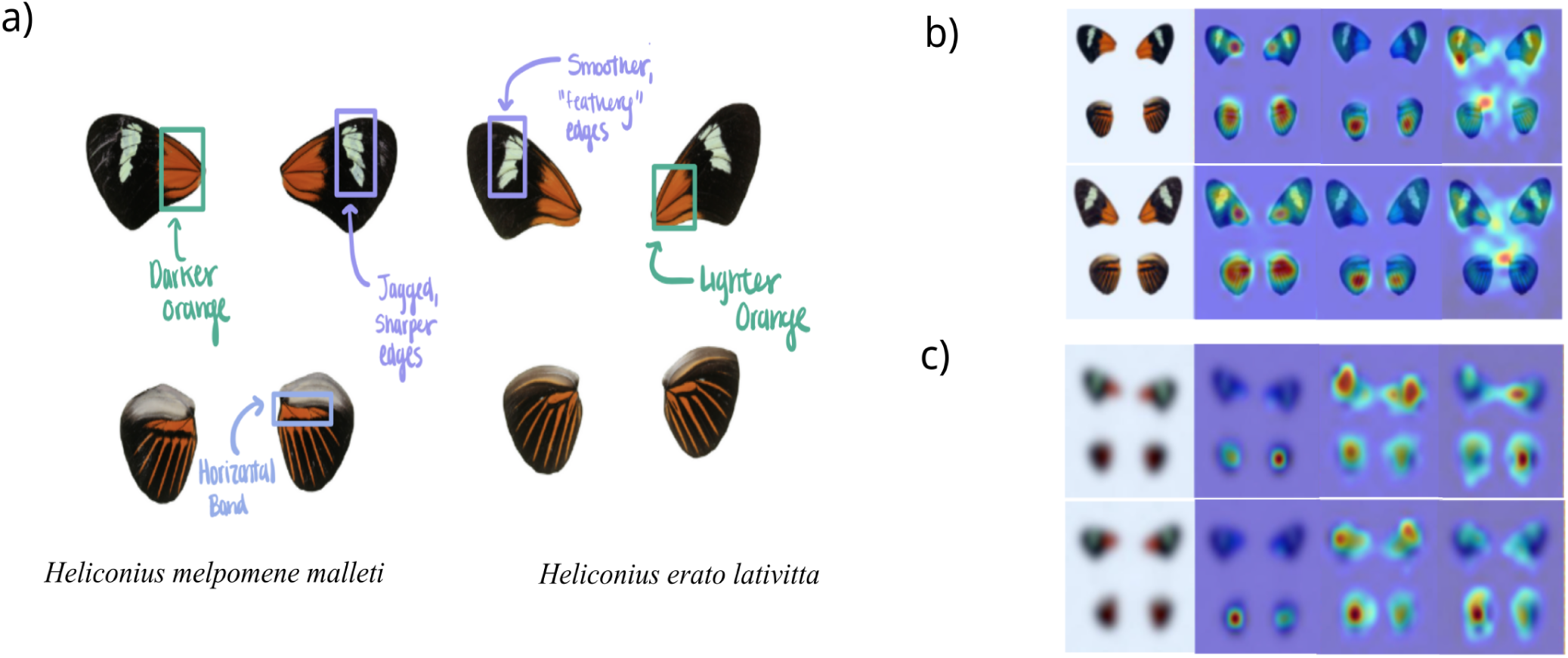
Diagnostic visual differences between mimetic subspecies of *Heliconius melpomene malleti* and *H. erato lativitta*. (a) Side-by-side comparison of mimetic subspecies, showing key diagnostic traits.(b, c) Model attention maps under simulated bird (b) and butterfly (c) visual acuity, using INTR.

Trait salience varied with acuity: avian-calibrated models emphasized localized features such as band curvature and patch boundaries (Fig. 6b), whereas butterfly-calibrated models prioritized broader wing regions (Fig. 6c), consistent with the lower spatial resolution of compound eyes. These findings suggest that the traits structuring mimetic similarity depend on the perceptual system of the observer.

## Discussion

The extraordinary diversity of *Heliconius* wing patterns has long been interpreted through the framework of discrete mimicry rings^9,10^. Our results instead indicate that mimicry is structured along perceptual continua rather than discrete categories, and shaped by both perception and geography. When phenotypes were quantified using high-dimensional neural network representations, traditional ring classifications were not consistently recovered as discrete clusters (Fig 2a). Forms differing in specific motifs—for example, the presence or absence of a hindwing yellow bar—often clustered closely in perceptual space, suggesting that certain pattern elements carry disproportionate weight in signal recognition^8,28^. Mimicry therefore appears to vary along a perceptual continuum rather than forming strictly bounded categories^26,29^.

Phenotypic similarity frequently did not reflect phylogenetic relatedness: both closely and distantly related species converged on similar color patterns. Phylogenetic signal was moderate but not statistically significant, indicating that shared ancestry explains only part of the observed variation (Fig 3a). This is consistent with previous work showing that genetic architecture can constrain, but does not fully determine, mimetic evolution^11,30^. In contrast, color pattern similarity aligned closely with geography (Fig 3b). Perceptual clusters corresponded to major biogeographic regions, including the Andes and adjacent lowlands ^31,32^, and boundaries in perceptual space matched documented hybrid zones^33–36^. The Andes emerged as a focal region of phenotypic diversification, consistent with their established role in *Heliconius* evolution^37,38^. Therefore, local ecological conditions organize mimetic similarity more strongly than shared ancestry alone.

Geographic overlap analyses further suggest that mimicry is embedded within community context rather than solely pairwise interactions^25,39^. Some co-mimetic pairs exhibited extensive niche overlap, consistent with strong local frequency-dependent selection requiring co-occurrence. Others shared similar phenotypes despite limited overlap (Fig. 4). This heterogeneity indicates that the spatial scale of frequency-dependent selection differs among mimicry systems^19,40,41^. In species-poor regions, co-occurrence may be necessary to maintain signal efficacy. In more diverse communities, multiple co-mimics may collectively reinforce the same warning signal, allowing phenotypic convergence to persist even when individual taxa rarely overlap. Mimicry thus appears to be structured not only by bilateral interactions, but by the broader composition of local communities^42^.

Deep learning approaches have previously demonstrated that mimetic convergence in *Heliconius* can be recovered in high-dimensional phenotypic space^11^. Our analyses extend this framework by embedding biologically calibrated visual filters and explicitly testing whether convergence implies reciprocal signal equivalence.

When models were trained across all taxa, avian-calibrated networks achieved the highest classification accuracy, indicating that mimetic forms remain separable in feature space despite convergence (Table 1). In contrast, models parameterized to butterfly vision—particularly *Heliconius* females—showed lower accuracy, consistent with the reduced spatial resolution of compound eyes^22^. Notably, male-based visual models performed better than female-based models, aligning with known sexual dimorphism in ommatidia number and the importance of visual cues in mate recognition^24,43^. This suggests that sexual selection and mimicry act simultaneously on wing pattern evolution, with male discrimination potentially maintaining within-species signal fidelity even as predator-mediated selection favors convergence ^44^.

When training was restricted to individual lineages, generalization was asymmetric. Models trained on *H. erato* transferred less effectively to *H. melpomene* than the reverse, a pattern consistent across visual regimes. Although differences in sample size and within-lineage variance may contribute (See Supplemental Figure 1), this asymmetry suggests that convergence may not always be reciprocal. If one lineage provides a more stable or more easily learned warning template, mimicry may be biased toward advergence (one species evolving towards the other) rather than balanced Müllerian mutualism^45^. Differences in demographic history, population structure, or relative abundance could further reinforce such asymmetries by influencing signal stability, predator learning dynamics, or even levels of toxicity^41,42,46^. For example, variation in demographic context has been shown to shift mimetic interactions in the monarch-viceroy mimicry system^47^. Mimicry rings may therefore contain internal asymmetries in their contribution to shared warning signals, reflecting dynamic rather than strictly reciprocal evolutionary interactions.

Variation in feature salience across visual models further informs the persistence of imperfect mimicry^7^. Avian-calibrated networks prioritized localized elements such as forewing bands and patches, whereas *Heliconius*-calibrated models integrated broader spatial regions (Figure 5b). Moreover, discrimination is constrained by perceptual thresholds: under Weber’s law, only differences exceeding a proportional just-noticeable threshold are likely to influence recognition^48^. For mobile avian predators viewing prey at variable distances and brief exposure times, fine-scale differences in elements such as hindwing ray width may fall below this threshold (Fig 5a). Imperfect mimicry may therefore persist not because selection is uniformly relaxed, but because certain trait differences fall below perceptual discrimination thresholds of relevant receivers.

Several limitations temper these inferences. Neural network classification does not directly model predator learning dynamics, decision thresholds or fitness consequences in natural populations, and biologically calibrated filters capture only a subset of perceptual dimensions^49^.

Ultraviolet reflectance, motion cues, behavioral context, viewing distance, environmental background complexity, and multi-modal integration were not incorporated^44,50–54^, all of which influence selection on color pattern. Because models were trained on standardized dorsal-view images, they do not capture variation in posture or ecological light environments that shape perception in the wild^44,55,56^. Asymmetric generalization may also be influenced by sampling differences or variance structure within lineages (See Supplemental Figure S1). Our analyses therefore quantify perceptual similarity under defined assumptions, but do not reconstruct predator learning or selection dynamics in natural contexts.

Together, these findings indicate that *Heliconius* mimicry is not a static system of discrete, reciprocally defined rings, but a receiver-dependent and geographically structured process in which convergence can be asymmetric and trait salience varies among observers. Integrating perceptual modeling with behavioral validation and ecological data will be essential for determining how similar is similar enough for protection. By linking perception, community structure and evolutionary history, machine learning provides a quantitative framework for testing how warning signals are structured, maintained and transformed within ecological communities.

## Methods

### Study System and Image Dataset

We restricted analyses to *Heliconius erato* and *H. melpomene*, the most extensively studied Müllerian mimicry pair, to test for asymmetry in mimetic resemblance. Focusing on this canonical species pair enabled direct comparison with Hoyal-Cuthill et al. (2019), who quantified mimetic similarity and explicitly assessed asymmetry in the same system^11^. Using the same focal taxa facilitates methodological continuity and comparability of results. Additional *Heliconius* mimicry rings were excluded because overlapping co-mimetic networks and distinct evolutionary histories could confound inference about symmetry in mimetic resemblance.

Standardized dorsal whole-body images of *Heliconius erato* and *Heliconius melpomene* subspecies were obtained from the Clinique Heliconius website (https://cliniquevetodax.com/Heliconius/index.html), which documents co-mimetic relationships. Images were resized to 1280 × 1280 pixels to preserve spatial resolution while minimizing computational load. Co-mimetic pairs were defined according to documented mimicry associations, and representative color pattern templates were constructed using the pavo^57^ package in R to reduce artefactual variation from wing wear, lighting, and positioning. The final dataset comprised 56 subspecies (27 H. erato, 29 H. melpomene), yielding 22 co-mimetic pairs. Phylogenetic analyses were restricted to these 44 paired subspecies to focus on evolutionary relationships among forms directly involved in co-mimetic associations.

### Quantifying Mimetic Similarity

Mimetic similarity was quantified using a feature-based image analysis approach implemented with a pre-trained deep convolutional neural network^12,58,59^. Unlike pixel-wise or color histogram methods, CNNs analyze images hierarchically, allowing different aspects of the phenotype—such as band position, color contrast, edge definition, and spatial configuration—to contribute unequally to similarity^49^. Each image was reduced to a quantitative representation of its visual structure, and pairwise Euclidean distances provided a continuous measure of mimetic similarity^60,61^, with smaller values indicating greater resemblance. For visualization only, representations were projected into two dimensions using t-SNE ^62^; all statistical analyses were conducted in the original multidimensional space.

To test whether co-mimetic pairs were more similar than expected by chance, we generated a null distribution by randomly sampling 1,000 heterospecific (H. erato–H. melpomene) pairs and comparing observed distances using a two-sided Wilcoxon rank-sum test.

### Phylogenetic Structure of Mimetic Similarity

We next assessed whether similarity reflects shared ancestry. A phenotypic similarity tree was constructed using UPGMA clustering based on perceptual distances and compared with a composite phylogeny derived from a previous phylogeny made of Heliconius, pruned to the 44 subspecies analysed here^14,63,64^. Topological correspondence was visualized using tanglegrams. Phylogenetic signal was quantified using Blomberg’s K^16^ and Pagel’s λ^17^ calculated for the two t-SNE dimensions treated as continuous traits, with significance assessed using 1,000 permutations. Because t-SNE preserves local similarity relationships in the original perceptual space, its axes provide a reduced representation of phenotypic structure suitable for comparative analyses. These analyses distinguish clustering driven by shared ancestry from convergence consistent with mimicry.

### Geographic Structure and Range Overlap

To examine geographic structure, we analyzed georeferenced occurrence records from the Heliconius Maps dataset^18^, matched to subspecies-level phenotypic data. Occurrences were color-coded by perceptual position to visualize geographic arrangement of mimetic forms. To identify regions of phenotypic diversity and transition, records were aggregated onto a 0.4° raster grid and averaged within cells to generate a map of mean perceptual phenotype. Geographic overlap among subspecies was quantified using kernel density estimation and Schoener’s D^65^.

Overlap values were calculated for all subspecies pairs and extracted for co-mimetic pairs to assess whether mimetic subspecies co-occur more strongly than other heterospecific combinations. Distribution estimates also allowed comparison of geographic breadth among mimicry rings.

### Modeling Mimicry Under Visual Acuity

Because Müllerian mimicry is mediated by receiver perception, mimetic similarity must be evaluated under biologically realistic visual constraints. Spatial resolution (visual acuity) limits discrimination of fine-scale pattern differences and may alter mimetic similarity ^66^. We therefore modeled similarity under multiple acuity regimes. We analysed 3,822 dorsal-view Heliconius images^67–86^ from the Cambridge Butterfly Collection^23^. Non-biological artifacts were removed using LepidopteraLens^87^, and images were resized for visual modeling. We then used AcuityView^20,88^ to apply spatial filtering simulating specified acuity levels. We generated multiple acuity-filtered datasets (see Supplementary Table S2), including estimates based on measured Heliconius visual acuity^24^ and avian acuity parameters from the belted kingfisher (*Megaceryle alcyon*), the closest available relative of jacamars—Heliconius butterfly predators^19,21,40^

### Learning-Based Tests of Mimicry and Asymmetry

To evaluate mimetic similarity and asymmetry under these visual systems, we trained supervised image-classification models on the acuity-filtered datasets. We implemented a ResNet-50^89^ convolutional neural network fine-tuned to classify subspecies and incorporated an ArcFace^90^ loss function to enhance separation among learned categories. This classification-based framework models predator-like signal categorization more directly than triplet-loss approaches^11^, which optimize relative similarity without explicitly modeling category boundaries. While convolutional neural networks do not replicate avian visual processing in detail, this framework provides a tractable approximation of perceptual categorization under defined visual constraints. Training and test sets were stratified by subspecies to ensure balanced representation across classes, and performance metrics were calculated on held-out images not used during model fitting. Overall classification accuracy was used to assess how well subspecies remained distinguishable under different visual systems.

To test for asymmetry in mimetic resemblance, we trained separate models on H. *erato* (EratoNet) and H. *melpomene* (MelpomeneNet) and evaluated each model on unseen co-mimetic subspecies from the other clade. Asymmetry was assessed using confusion rates, specifically measuring how often images of one species were misclassified as their co-mimic relative to overall classification error. If patterns learned from one species generalize more readily to its co-mimic than the reverse, misclassification rates will differ between directions. Learned image representations were additionally extracted to compute pairwise distances between subspecies under each acuity regime; these distances were used for visualization and summary of mimetic structure (See Supplemental Figure4-Figure6).

### Visualization of Learned Structure and Trait Importance

To identify the visual traits driving classification, we applied INTR^27^, which generates heatmaps highlighting image regions contributing most strongly to model predictions. A subset of six subspecies were analyzed under three acuity conditions: control (human vision), butterfly (Heliconius male vision), and bird (kingfisher vision). Comparing heatmaps across visual systems allowed us to assess whether convergence operates at the level of specific pattern elements and whether perceptual limits alter the traits defining a mimicry ring.

### Software and Reproducibility

Machine learning analyses were conducted in Python using PyTorch^91^. All other analyses were performed in R^92^. Data manipulation and visualization used the tidyverse^93^. Phylogenetic analyses were conducted using ape^94^and phytools^63^. Geospatial analyses used sf ^95^ and terra^96^. Randomization procedures were seeded to ensure reproducibility. Full package versions and implementation details are provided in the Supplementary Information.

## References

1. Ruxton, G. D., Allen, W. L., Sherratt, T. N. & Speed, M. P. Avoiding Attack: The Evolutionary Ecology of Crypsis, Aposematism, and Mimicry. (Oxford University Press, 2019).

2. Jiggins, C. & Lamas, G. The Ecology and Evolution of Heliconius Butterflies. (2016). doi:10.1093/acprof:oso/9780199566570.001.0001.

3. Müller, F. Ituna and Thyridia: A remarkable case of mimicry in butterflies. Trans. Entomol. Soc. Lond. xx–xxix (1879).

4. Mallet, J. & Barton, N. H. STRONG NATURAL SELECTION IN A WARNING-COLOR HYBRID ZONE. Evol. Int. J. Org. Evol. 43, 421–431 (1989).

5. Mallet, J. & Joron, M. Evolution of Diversity in Warning Color and Mimicry: Polymorphisms, Shifting Balance, and Speciation. Annu. Rev. Ecol. Evol. Syst. 30, 201–233 (1999).

6. Penney, H. D., Hassall, C., Skevington, J. H., Abbott, K. R. & Sherratt, T. N. A comparative analysis of the evolution of imperfect mimicry. Nature 483, 461–464 (2012).

7. Kikuchi, D. W. & Pfennig, D. W. Imperfect Mimicry and the Limits of Natural Selection. Q. Rev. Biol. 88, 297–315 (2013).

8. Rossato, D. O. et al. Subtle variation in size and shape of the whole forewing and the red band among co-mimics revealed by geometric morphometric analysis in *Heliconius* butterflies. Ecol. Evol. 8, 3280–3295 (2018).

9. Brown, K. S. The Biology of Heliconius and Related Genera. Annu. Rev. Entomol. 26, 427–457 (1981).

10. Mallet, J. & Gilbert Jr., L. E. Why are there so many mimicry rings? Correlations between habitat, behaviour and mimicry in*Heliconius*butterflies. Biol. J. Linn. Soc. 55, 159–180 (1995).

11. Hoyal Cuthill, J. F., Guttenberg, N., Ledger, S., Crowther, R. & Huertas, B. Deep learning on butterfly phenotypes tests evolution’s oldest mathematical model. Sci. Adv. 5, eaaw4967 (2019).

12. Zhang, R., Isola, P., Efros, A. A., Shechtman, E. & Wang, O. The Unreasonable Effectiveness of Deep Features as a Perceptual Metric. Preprint at 10.48550/arXiv.1801.03924 (2018).

13. Felsenstein, J. Phylogenies and the Comparative Method. Am. Nat. 125, 1–15 (1985).

14. Blomberg, S. P., Garland JR., T. & Ives, A. R. Testing for Phylogenetic Signal in Comparative Data: Behavioral Traits Are More Labile. Evolution 57, 717–745 (2003).

15. Pagel, M. Inferring the historical patterns of biological evolution. Nature 401, 877–884 (1999).

16. Rosser, N. & Mallet, J. Interactive maps for visualizing geographic distributions and phenotypes. Trop. Lepidoptera Res. 104–107 (2024).

17. Chouteau, M., Arias, M. & Joron, M. Warning signals are under positive frequency-dependent selection in nature. Proc. Natl. Acad. Sci. 113, 2164–2169 (2016).

18. Caves, E. M. & Johnsen, S. AcuityView: An r package for portraying the effects of visual acuity on scenes observed by an animal. Methods Ecol. Evol. 9, 793–797 (2018).

19. Caves, E. M., Fernández-Juricic, E. & Kelley, L. A. Ecological and morphological correlates of visual acuity in birds. J. Exp. Biol. 227, jeb246063 (2024).

20. Land, M. F. VISUAL ACUITY IN INSECTS. Annu. Rev. Entomol. 42, 147–177 (1997).

21. Jiggins, C., Warren, I., Salazar, P. & Montejo-Kovacevich, G. Heliconiine Butterfly Collection Records from University of Cambridge. (2025) doi:10.15468/qabb9f.

22. Wright, D. S. et al. Quantifying visual acuity in Heliconius butterflies. Biol. Lett. 19, 20230476 (2023).

23. Joron, M. & Iwasa, Y. The evolution of a Müllerian mimic in a spatially distributed community. J. Theor. Biol. 237, 87–103 (2005).

24. Balogh, A. C. V., Gamberale-Stille, G. & Leimar, O. Learning and the mimicry spectrum: from quasi-Bates to super-Müller. Anim. Behav. 76, 1591–1599 (2008).

25. Paul, D., et al. A Simple Interpretable Transformer for Fine-Grained Image Classification and Analysis. Preprint at 10.48550/arXiv.2311.04157 (2024).

26. Smith, S. H., Queste, L. M., Wright, D. S., Bacquet, C. N. & Merrill, R. M. Mating preferences act independently on different elements of visual signals in Heliconius butterflies. Behav. Ecol. 35, arae056 (2024).

27. Speed, M. P. & Turner, J. R. G. Learning and memory in mimicry: II. Do we understand the mimicry spectrum? Biol. J. Linn. Soc. 67, 281–312 (1999).

28. Van Belleghem, S. M., Alicea Roman, P. A., Carbia Gutierrez, H., Counterman, B. A. & Papa, R. Perfect mimicry between Heliconius butterflies is constrained by genetics and development. Proc. R. Soc. B Biol. Sci. 287, 20201267 (2020).

29. Jiggins, C. D. Ecological Speciation in Mimetic Butterflies. BioScience 58, 541–548 (2008).

30. Pérochon, E. et al. Müllerian Mimicry in Neotropical Butterflies: One Mimicry Ring to Bring Them All and in the Jungle Bind Them. Glob. Ecol. Biogeogr. 34, e70127 (2025).

31. Jiggins, C. D., McMillan, W. O., Neukirchen, W. & Mallet, J. What can hybrid zones tell us about speciation? The case of Heliconius erato and H. himera (Lepidoptera: Nymphalidae). Biol. J. Linn. Soc. 59, 221–242 (1996).

32. Blum, M. J. Ecological and genetic associations across a *Heliconius* hybrid zone. J. Evol. Biol. 21, 330–341 (2008).

33. Rosser, N., Dasmahapatra, K. K. & Mallet, J. Stable Heliconius butterfly hybrid zones are correlated with a local rainfall peak at the edge of the Amazon basin. Evolution 68, 3470–3484 (2014).

34. Pereira Martins, A. R., et al. Scale-dependent environmental effects on phenotypic distributions in Heliconius butterflies. Ecol. Evol. 12, e9286 (2022).

35. Rosser, N., Phillimore, A. B., Huertas, B., Willmott, K. R. & Mallet, J. Testing historical explanations for gradients in species richness in heliconiine butterflies of tropical America: DIVERSIFICATION OF BUTTERFLIES. Biol. J. Linn. Soc. 105, 479–497 (2012).

36. Rueda-M, N., Salgado-Roa, F. C., Gantiva-Q, C. H., Pardo-Díaz, C. & Salazar, C. Environmental Drivers of Diversification and Hybridization in Neotropical Butterflies. Front. Ecol. Evol. 9, (2021).

37. Elias, M., Gompert, Z., Jiggins, C. & Willmott, K. Mutualistic Interactions Drive Ecological Niche Convergence in a Diverse Butterfly Community. PLOS Biol. 6, e300 (2008).

38. Arias, M. et al. Crossing fitness valleys: empirical estimation of a fitness landscape associated with polymorphic mimicry. Proc. Biol. Sci. 283, 20160391 (2016).

39. Ogilvie, J. G. et al. Balanced polymorphisms and their divergence in a Heliconius butterfly. Ecol. Evol. 11, 18319–18330 (2021).

40. Willmott, K. R., Robinson Willmott, J. C., Elias, M. & Jiggins, C. D. Maintaining mimicry diversity: optimal warning colour patterns differ among microhabitats in Amazonian clearwing butterflies. Proc. R. Soc. B Biol. Sci. 284, 20170744 (2017).

41. McCulloch, K. J. et al. Sexual Dimorphism and Retinal Mosaic Diversification following the Evolution of a Violet Receptor in Butterflies. Mol. Biol. Evol. 34, 2271–2284 (2017).

42. Dell’Aglio, D. D., Troscianko, J., McMillan, W. O., Stevens, M. & Jiggins, C. D. The appearance of mimetic Heliconius butterflies to predators and conspecifics. Evolution 72, 2156–2166 (2018).

43. Mallet, J. Causes and Consequences of a Lack of Coevolution in M??llerian mimicry. Evol Ecol 13, 777–806 (1999).

44. Flanagan, N. S. et al. Historical demography of Müllerian mimicry in the neotropical *Heliconius* butterflies. Proc. Natl. Acad. Sci. 101, 9704–9709 (2004).

45. Prudic, K. L., Timmermann, B. N., Papaj, D. R., Ritland, D. B. & Oliver, J. C. Mimicry in viceroy butterflies is dependent on abundance of the model queen butterfly. *Commun*. Biol. 2, 68 (2019).

46. Dixit, T., Caves, E. M., Spottiswoode, C. N. & Horrocks, N. P. C. Why and how to apply Weber’s Law to coevolution and mimicry. Evol. Int. J. Org. Evol. 75, 1906–1919 (2021).

47. Lürig, M. D., Donoughe, S., Svensson, E. I., Porto, A. & Tsuboi, M. Computer Vision, Machine Learning, and the Promise of Phenomics in Ecology and Evolutionary Biology. Front. Ecol. Evol. 9, 642774 (2021).

48. Bybee, S. M. et al. UV photoreceptors and UV-yellow wing pigments in Heliconius butterflies allow a color signal to serve both mimicry and intraspecific communication. Am. Nat. 179, 38–51 (2012).

49. Page, E. et al. Pervasive mimicry in flight behavior among aposematic butterflies. Proc. Natl. Acad. Sci. 121, e2300886121 (2024).

50. Mérot, C., Frérot, B., Leppik, E. & Joron, M. Beyond magic traits: Multimodal mating cues in Heliconius butterflies. Evolution 69, 2891–2904 (2015).

51. Hausmann, A. E. et al. Light environment influences mating behaviours during the early stages of divergence in tropical butterflies. Proc. R. Soc. B Biol. Sci. 288, 20210157 (2021).

52. Montgomery, S. H., Rossi, M., McMillan, W. O. & Merrill, R. M. Neural divergence and hybrid disruption between ecologically isolated Heliconius butterflies. Proc. Natl. Acad. Sci. 118, e2015102118 (2021).

53. Dell’Aglio, D. D., Stevens, M. & Jiggins, C. D. Avoidance of an aposematically coloured butterfly by wild birds in a tropical forest. Ecol. Entomol. 41, 627–632 (2016).

54. Dell’Aglio, D. D., Troscianko, J., Stevens, M., McMillan, W. O. & Jiggins, C. D. The conspicuousness of the toxic *Heliconius* butterflies across time and habitat. Preprint at 10.1101/662155 (2019).

55. Maia, R., Gruson, H., Endler, J. A. & White, T. E. pavo 2: New tools for the spectral and spatial analysis of colour in r. Methods Ecol. Evol. 10, 1097–1107 (2019).

56. Deng, J. et al. ImageNet: A large-scale hierarchical image database. in 2009 IEEE Conference on Computer Vision and Pattern Recognition 248–255 (2009). doi:10.1109/CVPR.2009.5206848.

57. Aloysius, N. & Geetha, M. A review on deep convolutional neural networks. in 2017 International Conference on Communication and Signal Processing (ICCSP) 0588–0592 (2017). doi:10.1109/ICCSP.2017.8286426.

58. Wham, D. C., Ezray, B. & Hines, H. M. Measuring Perceptual Distance of Organismal Color Pattern using the Features of Deep Neural Networks. 736306 Preprint at 10.1101/736306 (2019).

59. Ezray, B. D., Wham, D. C., Hill, C. E. & Hines, H. M. Unsupervised machine learning reveals mimicry complexes in bumblebees occur along a perceptual continuum. Proc. R. Soc. B Biol. Sci. 286, 20191501 (2019).

60. van der Maaten, L. & Hinton, G. Visualizing Data using t-SNE. J. Mach. Learn. Res. 9, 2579–2605 (2008).

61. Kozak, K. M. et al. Multilocus Species Trees Show the Recent Adaptive Radiation of the Mimetic Heliconius Butterflies. Syst. Biol. 64, 505–524 (2015).

62. Revell, L. J. phytools: an R package for phylogenetic comparative biology (and other things). Methods Ecol. Evol. 3, 217–223 (2012).

63. Sneath, P. H. A. BASIC program for a significance test for clusters in UPGMA dendrograms obtained from squared Euclidean distances. Comput. Geosci. 5, 127–137 (1979).

64. Warren, D. L., Glor, R. E. & Turelli, M. Environmental niche equivalency versus conservatism: quantitative approaches to niche evolution. Evol. Int. J. Org. Evol. 62, 2868–2883 (2008).

65. Wilson, J. S., Jahner, J. P., Williams, K. A. & Forister, M. L. Ecological and Evolutionary Processes Drive the Origin and Maintenance of Imperfect Mimicry. PLoS ONE 8, e61610 (2013).

66. Lawrence, C. & Campolongo, E. G. Heliconius Collection (Cambridge Butterfly). (2024) doi:10.57967/hf/2668.

67. Montejo-Kovacevich, G., van der Heijden, E., Nadeau, N. & Jiggins, C. Cambridge butterfly wing collection batch 10. (2020) doi:10.5281/zenodo.4289223.

68. Salazar, P. A., Nadeau, N., Montejo-Kovacevich, G. & Jiggins, C. Sheffield butterfly wing collection - Patricio Salazar, Nicola Nadeau, Ikiam broods batch 1 and 2. (2020) doi:10.5281/zenodo.4288311.

69. Jiggins, C. & Warren, I. Cambridge butterfly wing collection - Chris Jiggins 2001/2 broods batch 2. (2019) doi:10.5281/zenodo.2550097.

70. Jiggins, C., Montejo-Kovacevich, G., Warren, I. & Wiltshire, E. Cambridge butterfly wing collection batch 3. (2019) doi:10.5281/zenodo.2682458.

71. Montejo-Kovacevich, G., Jiggins, C. & Warren, I. Cambridge butterfly wing collection batch 4. (2019) doi:10.5281/zenodo.2682669.

72. Montejo-Kovacevich, G., Jiggins, C., Warren, I. & Wiltshire, E. Cambridge butterfly wing collection batch 5. (2019) doi:10.5281/zenodo.2684906.

73. Warren, I. & Jiggins, C. Miscellaneous Heliconius wing photographs (2001-2019) Part 1. (2019) doi:10.5281/zenodo.2552371.

74. Warren, I. & Jiggins, C. Miscellaneous Heliconius wing photographs (2001-2019) Part 3. (2019) doi:10.5281/zenodo.2553977.

75. Montejo-Kovacevich, G., Jiggins, C., Warren, I. & Wiltshire, E. Cambridge butterfly wing collection batch 6. (2019) doi:10.5281/zenodo.2686762.

76. Jiggins, C. & Warren, I. Cambridge butterfly wing collection - Chris Jiggins 2001/2 broods batch 1. (2019) doi:10.5281/zenodo.2549524.

77. Meier, J. I., Salazar, P., Montejo-Kovacevich, G., Warren, I. & Jggins, C. Cambridge butterfly wing collection - Patricio Salazar PhD wild specimens batch 3. (2020) doi:10.5281/zenodo.4153502.

78. Montejo-Kovacevich, G., Jiggins, C. & Warren, I. Cambridge butterfly wing collection batch 1-version 2. (2019) doi:10.5281/zenodo.3082688.

79. Salazar, C., Montejo-Kovacevich, G., Jiggins, C., Warren, I. & Gavins, I. Camilo Salazar and Cambridge butterfly wing collection batch 1. (2019) doi:10.5281/zenodo.2735056.

80. Montejo-Kovacevich, G., Jiggins, C., Warren, I. & Wiltshire, E. Cambridge butterfly wing collection batch 7. (2019) doi:10.5281/zenodo.2702457.

81. Pinheiro de Castro, E., et al. Brazilian Butterflies Collected December 2020 to January 2021. (2022) doi:10.5281/zenodo.5561246.

82. Montejo-Kovacevich, G., Jiggins, C., Warren, I. & Wiltshire, E. Cambridge butterfly wing collection batch 8. (2019) doi:10.5281/zenodo.2707828.

83. Montejo-Kovacevich, G., Jiggins, C., Warren, I., Wiltshire, E. & Gavins, I. Cambridge butterfly wing collection batch 9. (2019) doi:10.5281/zenodo.2714333.

84. Montejo-Kovacevich, G., van der Heijden, E. & Jiggins, C. Cambridge butterfly collection - GMK Broods Ikiam 2018. (2020) doi:10.5281/zenodo.4291095.

85. Montejo-Kovacevich, G. et al. Cambridge and collaborators butterfly wing collection batch 10. (2019) doi:10.5281/zenodo.2813153.

86. Ramirez, M. Lepidoptera Wing Segmentation. 10.5281/zenodo.10869579 (2024).

87. van den Berg, C. P., Troscianko, J., Endler, J. A., Marshall, N. J. & Cheney, K. L. Quantitative Colour Pattern Analysis (QCPA): A comprehensive framework for the analysis of colour patterns in nature. Methods Ecol. Evol. 11, 316–332 (2020).

88. He, K., Zhang, X., Ren, S. & Sun, J. Deep Residual Learning for Image Recognition. Preprint at 10.48550/arXiv.1512.03385 (2015).

89. Deng, J. et al. ArcFace: Additive Angular Margin Loss for Deep Face Recognition. IEEE Trans. Pattern Anal. Mach. Intell. 44, 5962–5979 (2022).

90. Paszke, A., et al. PyTorch: An Imperative Style, High-Performance Deep Learning Library. Preprint at 10.48550/arXiv.1912.01703 (2019).

91. R Core Team. R: A Language and Environment for Statistical Computing. (R Foundation for Statistical Computing, Vienna, Austria, 2021).

92. Wickham, H. et al. Welcome to the tidyverse. J. Open Source Softw. 4, 1686 (2019).

93. Paradis, E. & Schliep, K. ape 5.0: an environment for modern phylogenetics and evolutionary analyses in R. Bioinformatics 35, 526–528 (2019).

94. Pebesma, E. Simple Features for R: Standardized Support for Spatial Vector Data. R J. 10, 439–446 (2018).

95. Hijmans, R. J., Brown, A. & Barbosa, M. Terra: Spatial Data Analysis. (2026).

