## Supplemental Materials for "The perceptual and spatial architecture of Müllerian mimicry in *Heliconius* Butterflies"

This PDF file includes:

Figure S1. Class distribution across training, validation and test datasets for subspecies classification.
Figure S2.  Permutation test of mimetic phenotypic similarity.
Table S1. Mimetic subspecies pairs included in classification-based modeling analyses.
Table S2. Visual acuity parameters used for perceptual filtering in AcuityView.
Table S3. Classification accuracy across visual acuity regimes and training frameworks.
Figure S3. Perceptual distance distributions across visual acuity regimes and training frameworks.
Figure S4–S6.  t-SNE visualizations of perceptual embeddings under alternative visual acuity regimes.
Figure S7. Subtle pattern differences among representative co-mimetic subspecies pairs.
Figure S8. Model attention maps under alternative visual acuity regimes.

Other Supplemental Material for this manuscript includes the following:

Lawrence, C. (2026). The perceptual and spatial architecture of Müllerian mimicry in Heliconius Butterflies (Zenodo).  https://doi.org/10.5281/zenodo.18838627

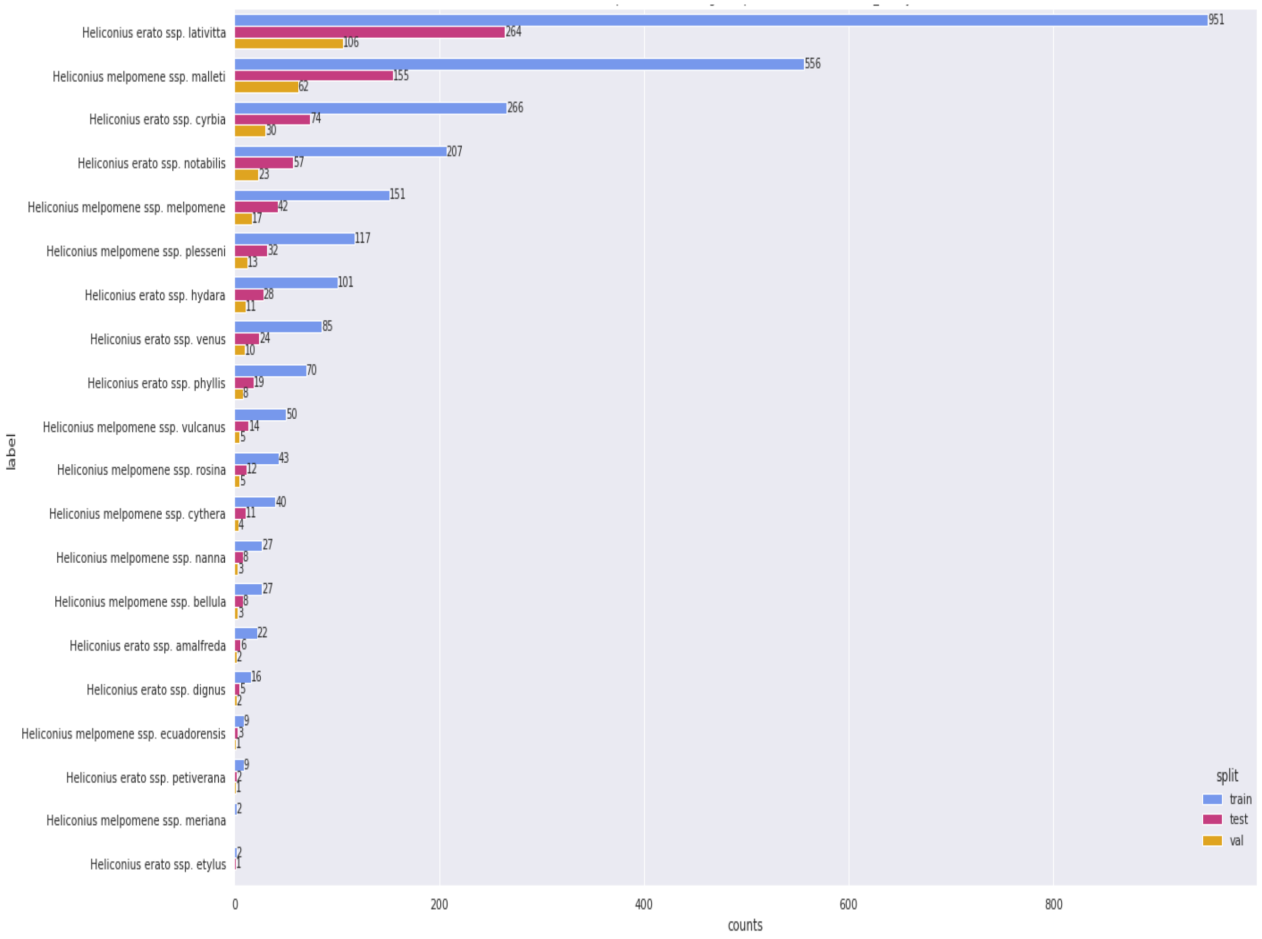


**Figure S1.** Class distribution across training, validation and test datasets for subspecies classification. Distribution of image counts per subspecies of *Heliconius erato* and *H. melpomene* used to train and evaluate the baseline classification model prior to application of biologically calibrated visual filters. Bars indicate the number of images assigned to the training (blue), validation (gold) and test (red) partitions for each subspecies. The dataset is unbalanced, with greater overall representation of *H. erato* relative to *H. melpomene*, and particularly high sampling of *H. erato lativitta*. This skew in class frequency motivated the reporting of both microaccuracy (overall classification rate) and macroaccuracy (mean per-class accuracy) in subsequent analyses, as macroaccuracy mitigates the influence of uneven sampling across taxa.

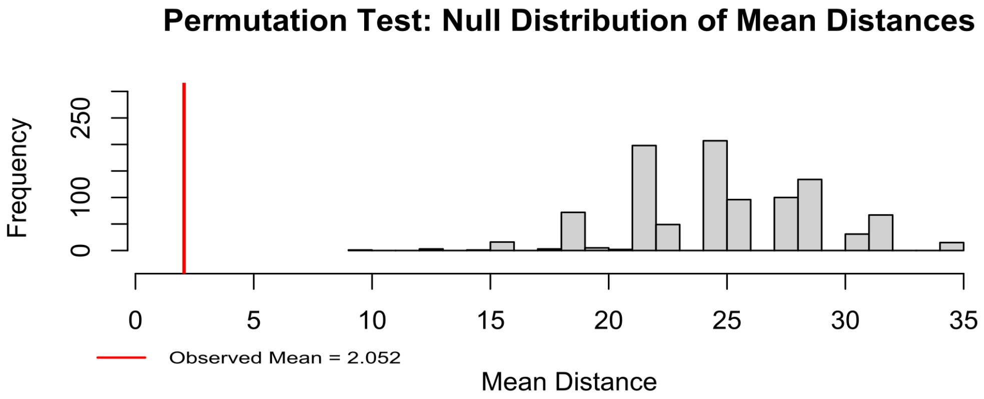


**Figure S2.** Permutation test of mimetic phenotypic similarity.
Permutation-based assessment of whether empirically defined mimicry pairs exhibit greater phenotypic similarity than expected by chance. The observed mean Euclidean distance among known mimetic pairs in perceptual (t-SNE) space was 2.051. To generate a null expectation, 1,000 permuted datasets were constructed by randomly sampling species without replacement from the full pool and forming the same number of interspecific pairs as in the empirical dataset (n = 28 pairs per permutation). For each permutation, the mean pairwise distance was calculated, yielding a null distribution of mean distances under random pairing. The observed mean distance (vertical line) lies in the extreme lower tail of the null distribution. Only one of 1,001 total values (including the observed value) was equal to or smaller than the empirical mean, corresponding to an empirical one-tailed p-value of 0.001. These results indicate that co-mimetic taxa are significantly more phenotypically similar than expected under random species associations.

| Mimic Pairs | |
|---|---|
| **Heliconius melpomene** | **Heliconius erato** |
| malleti | lativitta |
| melpomene melpomene | hydara |
| plesseni | notabilis |
| vulcanus | venus |
| rosina | petiverana |
| cythera | cyrbia |
| nanna | phyllis |
| bellula | dignus |
| ecuadorensis | etylus |
| meriana | amalfreda |

**Table 1:** Mimic Pairs

**Table S1.** Mimetic subspecies pairs included in classification-based modeling analyses.
List of empirically recognized co-mimetic subspecies pairs of *Heliconius erato* and *H. melpomene* included in the supervised classification and cross-lineage generalization analyses. Pairs were defined based on documented mimicry associations and represent geographically corresponding warning-pattern forms. These pairings were used to (i) quantify cross-lineage perceptual similarity under biologically calibrated visual regimes and (ii) evaluate directional asymmetry in generalization between lineage-specific models (EratoNet and MelpomeneNet). Only subspecies for which standardized dorsal images and reliable taxonomic assignments were available were retained for modeling.

| Acuity | 1/cpd | Degrees |
|---|---|---|
| Male Behavioral Acuity | $\frac{1}{0.547}$ | 1.828 |
| Male Morphological Acuity | $\frac{1}{0.386}$ | 2.591 |
| Female Behavioral Acuity | $\frac{1}{0.428}$ | 2.336 |
| Female Morphological Acuity | $\frac{1}{0.369}$ | 2.710 |
| Kingfisher Acuity | $\frac{1}{26.0}$ | 0.038 |

**Table 2:** Acuity values used for image processing.

**Table S2.** Visual acuity parameters used for perceptual filtering in AcuityView.
Summary of spatial acuity parameters implemented to simulate biologically calibrated visual systems in AcuityView. For each visual regime, the estimated spatial resolution is reported in 1/cycles per degree (1/cpd).The final column lists the angular degree values applied within AcuityView to generate acuity-filtered images for each test condition. These parameters were derived from published estimates of visual acuity for Heliconius butterflies (male and female) and avian predators and were used to produce spatially filtered datasets for subsequent classification analyses.

| Acuity Condition | Model | Test On | Micro Acc | Macro Acc |
|---|---|---|---|---|
| No Acuity | AllNet | - | 0.963 | 0.838 |
| | EratoNet | Erato | 0.978 | 0.844 |
| | EratoNet | Melpomene | 0.971 | 0.771 |
| | MelpomeneNet | Melpomene | 0.965 | 0.892 |
| | MelpomeneNet | Erato | 0.955 | 0.869 |
| Heliconius Male Behavioral | AllNet | - | 0.939 | 0.804 |
| | EratoNet | Erato | 0.969 | 0.784 |
| | EratoNet | Melpomene | 0.945 | 0.704 |
| | MelpomeneNet | Melpomene | 0.965 | 0.927 |
| | MelpomeneNet | Erato | 0.971 | 0.913 |
| Heliconius Female Behavioral | AllNet | - | 0.915 | 0.767 |
| | EratoNet | Erato | 0.978 | 0.844 |
| | EratoNet | Melpomene | 0.933 | 0.711 |
| | MelpomeneNet | Melpomene | 0.949 | 0.904 |
| | MelpomeneNet | Erato | 0.945 | 0.870 |
| Heliconius Male Morphological | AllNet | - | 0.916 | 0.760 |
| | EratoNet | Erato | 0.983 | 0.873 |
| | EratoNet | Melpomene | 0.938 | 0.746 |
| | MelpomeneNet | Melpomene | 0.953 | 0.863 |
| | MelpomeneNet | Erato | 0.941 | 0.805 |
| Heliconius Female Morphological | AllNet | - | 0.898 | 0.732 |
| | EratoNet | Erato | 0.971 | 0.809 |
| | EratoNet | Melpomene | 0.943 | 0.751 |
| | MelpomeneNet | Melpomene | 0.957 | 0.859 |
| | MelpomeneNet | Erato | 0.963 | 0.853 |
| Kingfisher | AllNet | - | 0.972 | 0.875 |
| | EratoNet | Erato | 0.978 | 0.844 |
| | EratoNet | Melpomene | 0.964 | 0.733 |
| | MelpomeneNet | Melpomene | 0.973 | 0.898 |
| | MelpomeneNet | Erato | 0.955 | 0.847 |

| Model | Species seen at training time |
|---|---|
| AllNet | *H. melpomene* and *H. erato* |
| EratoNet | *H. erato* only |
| MelpomeneNet | *H. melpomene* |

**Table S3.** Classification accuracy across visual acuity regimes and training frameworks. Summary of classification performance under six biologically calibrated visual acuity conditions. The upper table reports microaccuracy (overall proportion of correctly classified images) and macroaccuracy (mean per-class accuracy) for each acuity regime and evaluation context/ The lower table lists each model and the species included during training, clarifying lineage-specific versus joint training configurations.

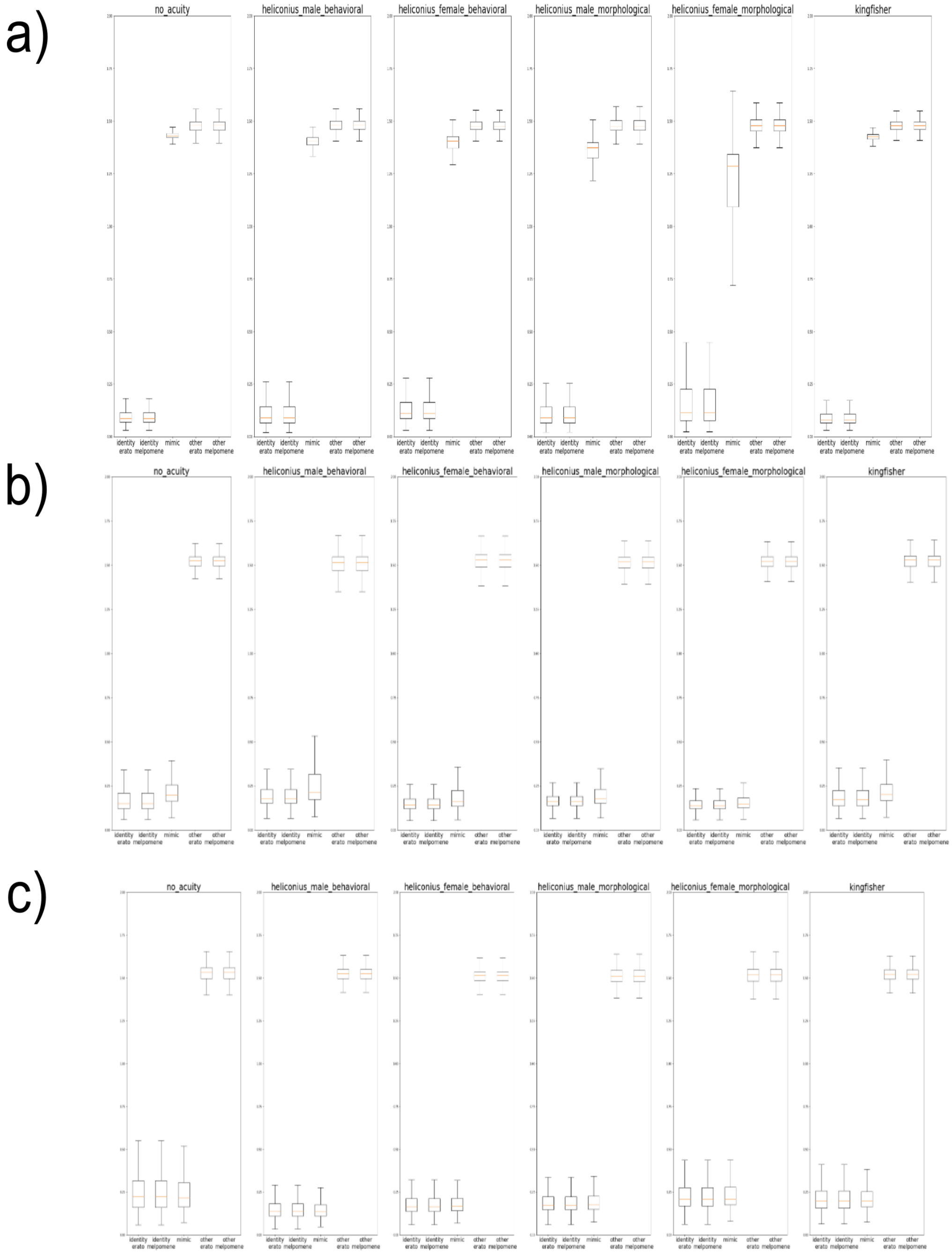


**Figure S3.** Perceptual distance distributions across visual acuity regimes and training frameworks. Boxplots of pairwise Euclidean distances in learned feature space under six biologically calibrated visual acuity regimes. Distances are grouped by outcome category: correct identification of *H. erato* ("identify erato"), correct identification of *H. melpomene* ("identify melpomene"), misclassification as a co-mimic ("mimic"), and misclassification to non-mimetic subspecies within *H. erato* ("other erato") or *H. melpomene* ("other melpomene"). Higher values indicate greater separation in feature space, whereas lower values indicate greater overlap. (a) **AllNet**, trained on both lineages.
(b) **EratoNet**, trained on *H. erato* only. (c) **MelpomeneNet**, trained on *H. melpomene* only.

**Figure S4–S6.** t-SNE visualizations of perceptual embeddings under alternative visual acuity regimes. t-distributed stochastic neighbor embedding (t-SNE) projections of learned feature representations for mimetic subspecies of *Heliconius erato* (lighter points) and *H. melpomene* (darker points). Each panel represents one of six biologically calibrated visual acuity conditions applied prior to model training. Points corresponding to co-mimetic subspecies pairs share color (distinct hues denote different mimicry pairs), allowing visualization of cross-lineage proximity in feature space.

Figure S4 shows embeddings from **AllNet** (trained jointly on both lineages), Figure S5 rom **EratoNet** (trained on *H. erato* only), and Figure S6 from **MelpomeneNet** (trained on *H. melpomene* only). All projections are derived from high-dimensional learned representations; distances in the original feature space were used for quantitative analyses.

**Fig S4.**

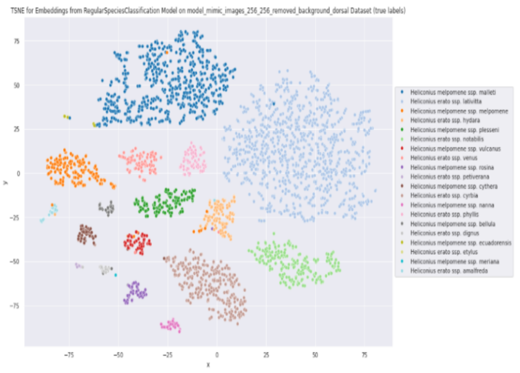

(a) Original

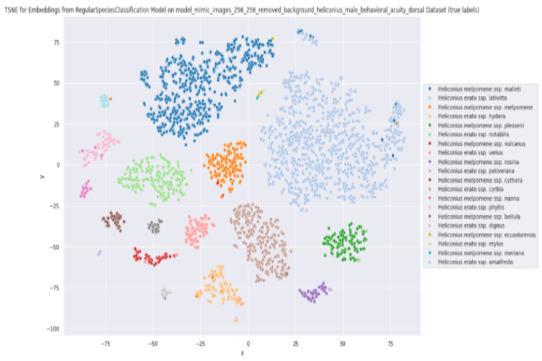

(b) Heliconius Male Behavioral Acuity

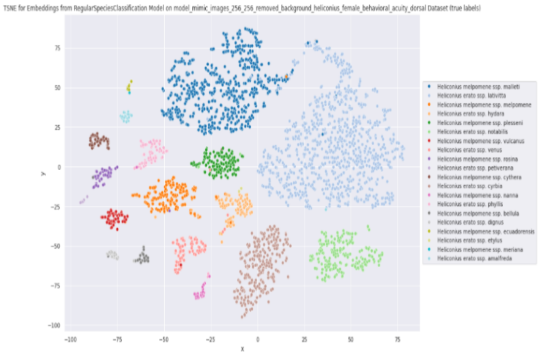

(c) Heliconius Female Behavioral Acuity

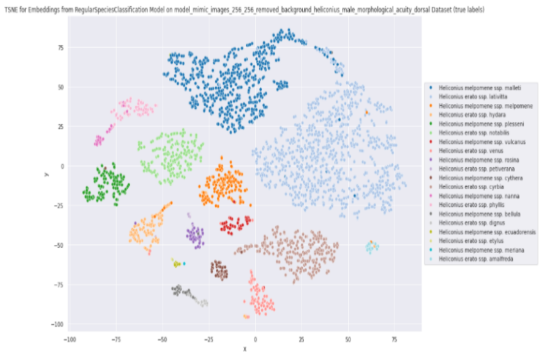

(d) Heliconius Male Morphological Acuity

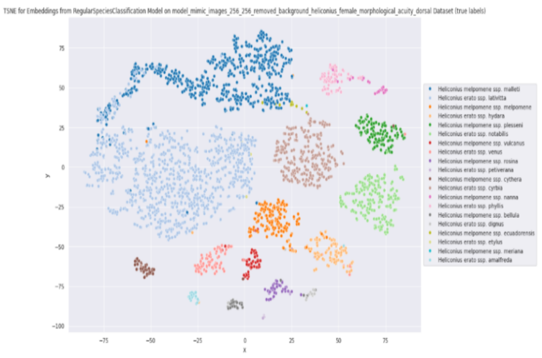

(e) Heliconius Female Morphological Acuity

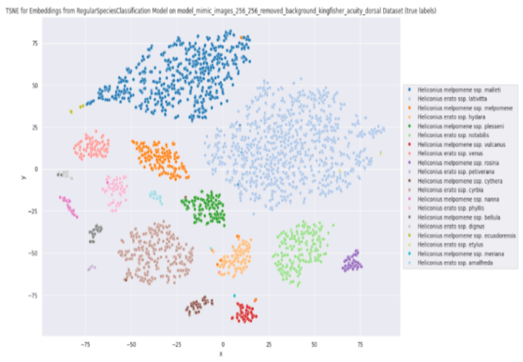

(f) Kingfisher Acuity

**Fig S5.**

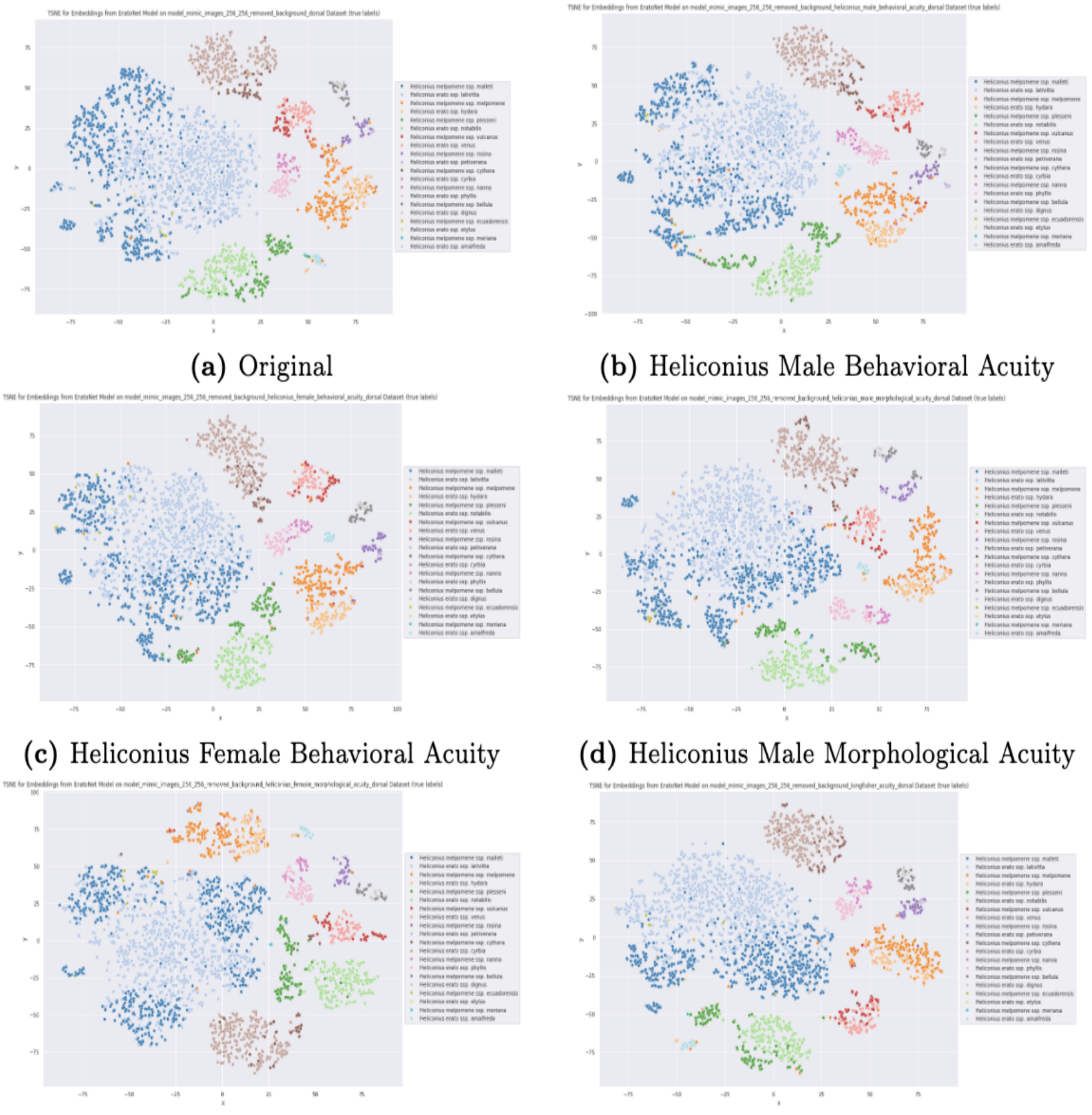

(a) Original
(b) Heliconius Male Behavioral Acuity
(c) Heliconius Female Behavioral Acuity
(d) Heliconius Male Morphological Acuity
(e) Heliconius Female Morphological Acuity
(f) Kingfisher Acuity

**Fig S6.**

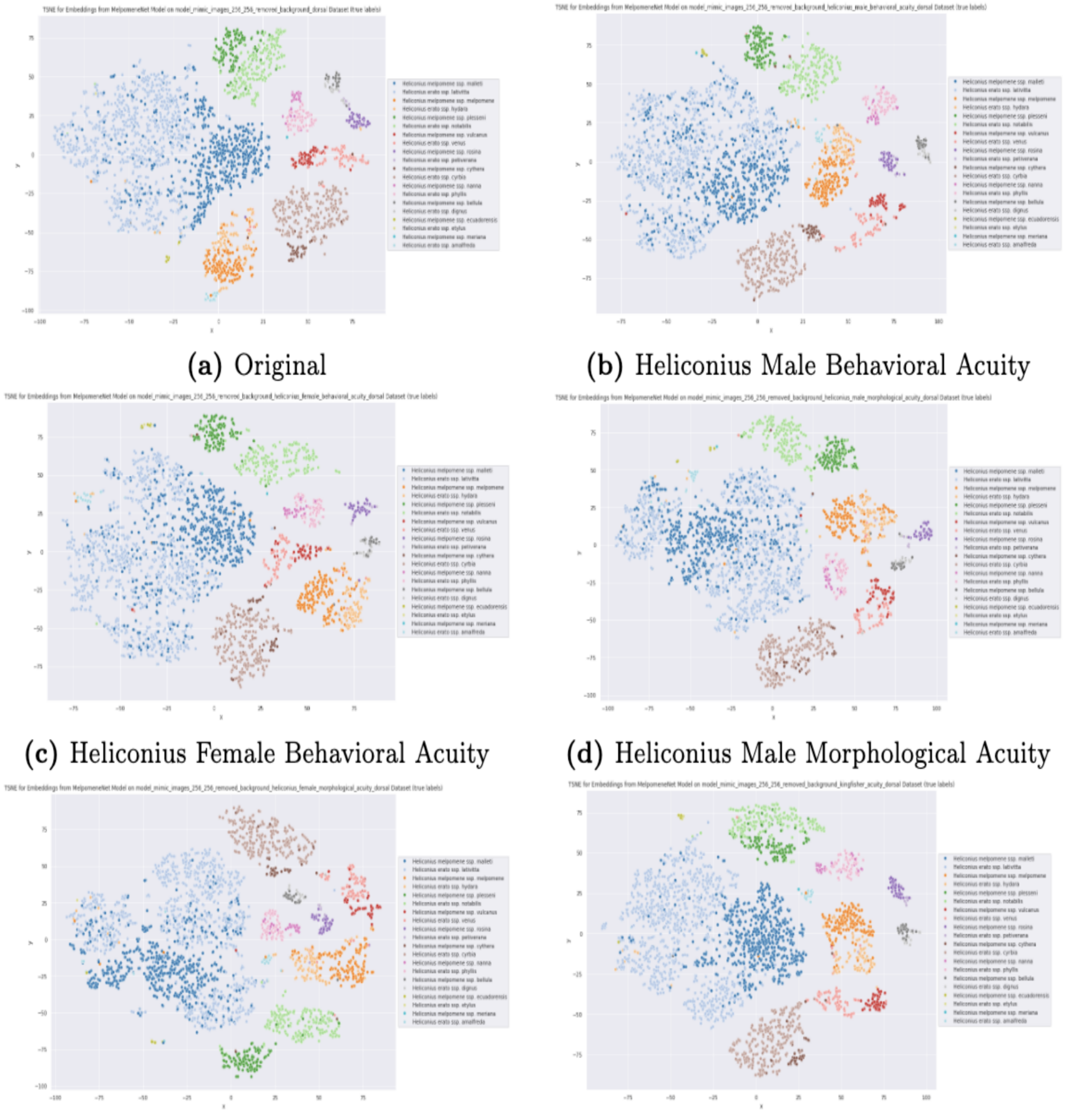

(a) Original

(b) Heliconius Male Behavioral Acuity

(c) Heliconius Female Behavioral Acuity

(d) Heliconius Male Morphological Acuity

(e) Heliconius Female Morphological Acuity

(f) Kingfisher Acuity

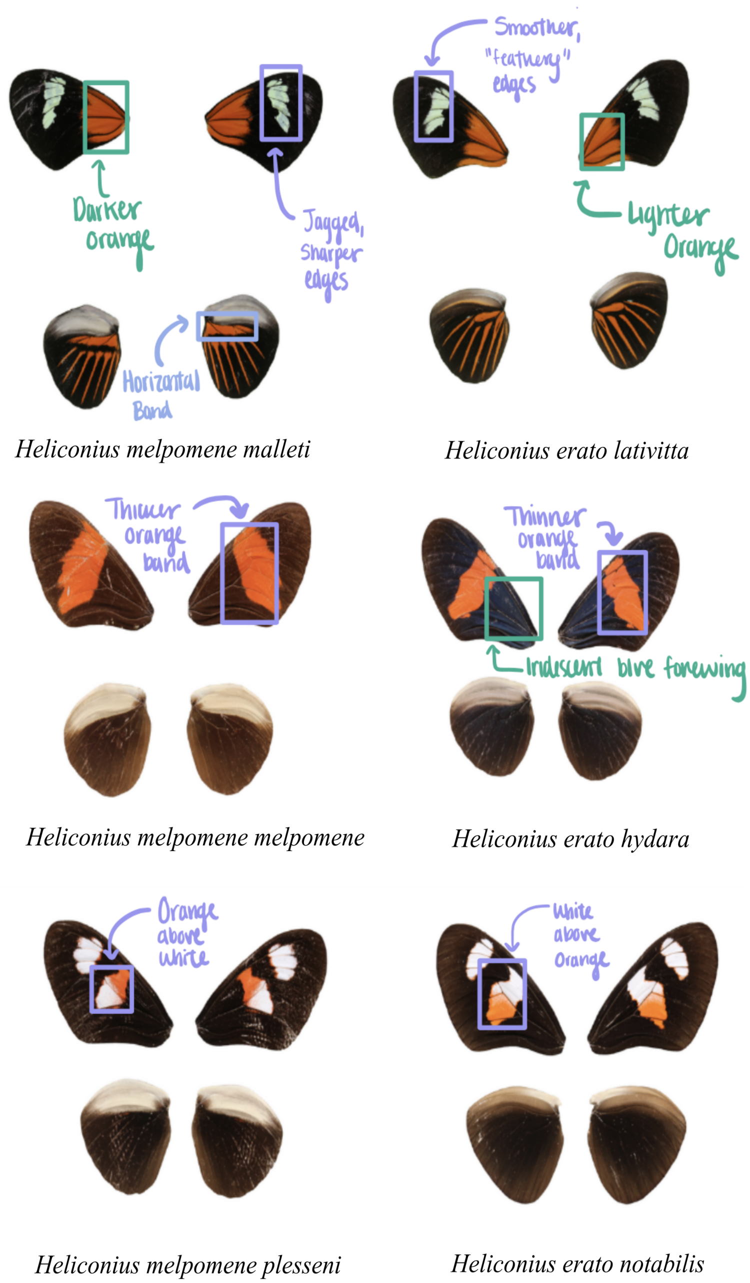


**Figure S7.** Subtle pattern differences among representative co-mimetic subspecies pairs. Representative dorsal wing patterns of three co-mimetic pairs of *Heliconius erato* and *H. melpomene*, highlighting subtle morphological differences within shared warning phenotypes. Shown are *H. melpomene malleti* and *H. erato lativitta*; *H. melpomene melpomene* and *H. erato hydara*; and *H. melpomene plesseni* and *H. erato notabilis*. Arrows indicate localized differences in forewing band shape, hindwing banding, and color patch boundaries between co-mimetic subspecies. These examples illustrate that phenotypes classified within the same mimicry association retain fine-scale pattern variation despite overall resemblance.

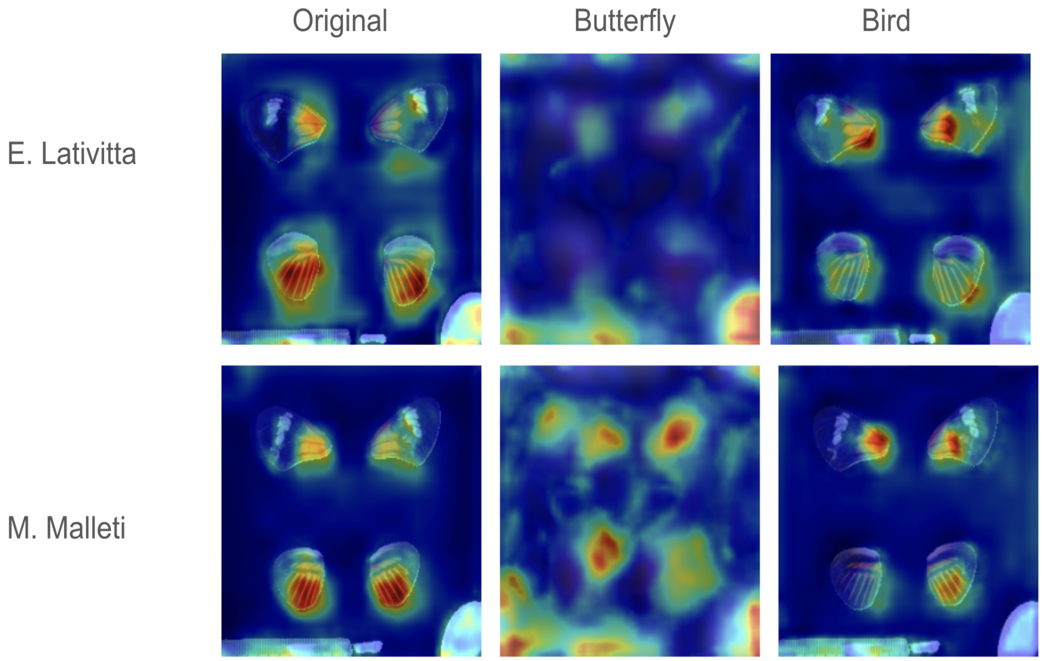


**Figure S8.** Model attention maps under alternative visual acuity regimes. Attention maps generated using INTR for co-mimetic subspecies *Heliconius erato lativitta* and *H. melpomene malleti* under three visual conditions. Heatmaps indicate image regions contributing most strongly to model predictions, with warmer colors denoting greater influence on classification decisions. Comparisons across visual regimes illustrate how spatial filtering alters the distribution of salient features identified by the model when discriminating between closely resembling mimetic forms.
